# SuperCT: A supervised-learning-framework to enhance the characterization of single-cell transcriptomic profiles

**DOI:** 10.1101/416719

**Authors:** Peng Xie, Mingxuan Gao, Chunming Wang, Pawan Noel, Chaoyong Yang, Daniel Von Hoff, Haiyong Han, Michael Q. Zhang, Wei Lin

**Affiliations:** University of Texas at Dallas. Department of Molecular and Cell Biology, Richardson, TX, USA; Xiamen University, Department of Chemical Biology, Xiamen, Fujian, China; National Engineering Research Center for Miniaturized Detection System, Chinese Northwest University, Xi’an, Shaanxi, China; Translational Genomics Research Institute, Molecular Medicine Division, Phoenix, AZ, USA

**Author notes:** Contribute equally (Peng Xie, Mingxuan Gao).

## Abstract

Characterization of individual cell types is fundamental to the study of multicellular samples such as tumor tissues. Single-cell RNAseq techniques, which allow high-throughput expression profiling of individual cells, have significantly advanced our ability of this task. Currently, most of the scRNA-seq data analyses are commenced with unsupervised clustering of cells followed by visualization of clusters in a low-dimensional space. Clusters are often assigned to different cell types based on canonical markers. However, the efficiency of characterizing the known cell types in this way is low and limited by the investigator[s] knowledge. In this study, we present a technical framework of training the expandable supervised-classifier in order to reveal the single-cell identities based on their RNA expression profiles. Using multiple scRNA-seq datasets we demonstrate the superior accuracy, robustness, compatibility and expandability of this new solution compared to the traditional methods. We use two examples of model upgrade to demonstrate how the projected evolution of the cell-type classifier is realized.

## Introduction

Categorizing individual cells into different cell types is an important step in characterizing complex tissues such as tumors. Recent advances in single-cell RNA-seq (scRNA-seq) techniques make it possible to profile the RNA transcript abundance in a single cell, which enables us to unveil its identity. The mainstream scRNA-seq analytical methods utilize dimensional reduction (DR) and unsupervised clustering (UC) algorithms to initiate the analyses. UC provides the mathematical aggregation based on some cell grouping measures and DR facilitates the visualization of the clustering result by “projection”. The putative subpopulations of cell types are thus identified with the enriched canonical signature signals. Nonetheless, this workflow has some limitations. First, the cell types were actually not characterized at the single-cell level but at the cluster level. The UC can take different distance/similarity metric that results in different clustering effects. Different people have the preference of metric implementing their own algorithms [1-4]. In addition, clustering is a so-called ill-defined problem in mathematics. Therefore, the interpretation of the resultant clustering could be arbitrary and method-dependent. Second, UC method highly relies on the investigator’s expert knowledge of the related cell markers or the functional molecules for result interpretation. Without such knowledge, the resultant clusters will be very hard to explain and some minor but critical populations could be easily overlooked. In this study, we sought to develop a new strategy that enables tackling the abovementioned problems. Moreover, we sought to rebuild a single-cell analytical framework that is not restricted by an individual investigator’s expertise, but expandable as the entire human knowledge of the living cells continuously grows.

Supervised classifier (SC) has been widely used in the automatic image classification [5-7]. Rämö *et. al.* developed CellClassifier based on the pixel intensities of cell imaging [8]. However, using only morphological information is obviously insufficient to find a definite answer. Characterizing single-cell gene expression profiles resembles the image recognition in terms of high-dimensional data transformation and classification. The transcriptome-wide RNA profiling of thousands of genes should be more than enough to discern a cell type from others even at relatively high noise level. All these facts make SC not only an alternative solution of scRNA-seq data analysis but also a more efficient one. In this study, we incorporate the Mouse-Cell-Atlas (MCA) datasets [9] for most of the training task of an artificial neural network (ANN) model. Using general principle of machine learning, we have built a robust SUPERvised Cell-Type classifier framework, called SuperCT to automatically learn the molecular features from the scRNA-seq data generated and classified by other investigators. This ANN-based automatic classifier can efficiently identify the similar cells in new samples. Moreover, using the strategy of online learning, the ANN model can continuously optimize the performance and adapt itself to the prediction tasks in a specific sample context using the training dataset generated from the similar background. By modifying the output structure and applying the transfer learning strategy, we are able to efficiently expand the cell-type catalogue for a broader scope of characterization task. In the current study, we extensively investigate the utility, the reliability, the compatibility and the expandability of the SuperCT framework and demonstrate with explicit examples on how to characterize cell types and gain unprecedented insight using this special technique.

## Result

The schematic workflow of SuperCT framework is shown in Fig. 1a. The detailed structure of the fully connected ANNs for SuperCT is described in the Online Methods. The training datasets were selected from MCA projct (∼190k cells) and peripheral blood mononuclear cells (PBMCs, ∼9k cells). More details of the training data organization are given in the Online Methods and **Supplementary Table 1.** The overall concordance measure is defined to assess the reliability of the prediction. More precisely, concordance means whether the SuperCT predictions agree with the manual annotation based on the enriched canonical marker of the entire cluster in a certain sample or the entire cellular sample sorted by the canonical markers from the specific tissue (there are a few cell types defined in this way in the MCA dataset).

**Figure 1:**
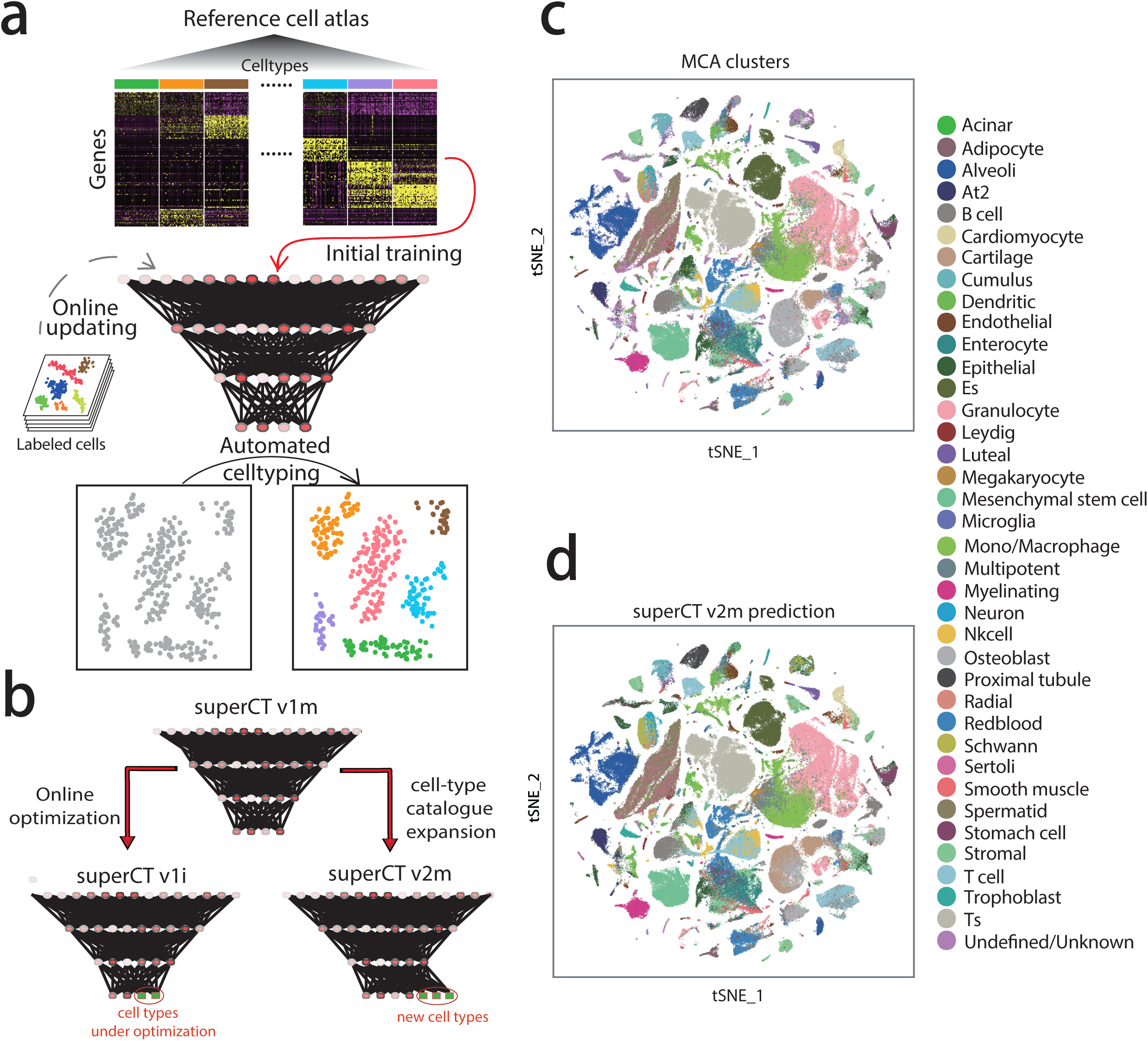
SuperCT framework and the high concordance between the original MCA cell-type annotations and the SuperCT predictions. a) The workflow of SuperCT training, prediction and upgrade. b) The two upgrades that lead to the optimized or expanded SuperCT classifier. c) & d) The overview of MCA data original labels in comparison with SuperCT v2m predictions.

Using a 2.3GHz CPU processor computer, SuperCT is able to process ∼10,000 single-cell expression profiles in less than one minute, whose efficiency is far superior to traditional methods that requires a lot of interactive operation and domain knowledge as input.

Other than the training dataset, in order to further validate the utility of SuperCT classifier in assigning cell types, we test its performances using three independent datasets. The first dataset was from peripheral blood mononuclear cells (PBMC). Stoeckius *et. al.* used a unique CITE-seq technique that allows to interrogate the single-cell transcriptomics and surface proteins at the same time [10]. The cell identities revealed by CITE-seq technique are validated by both RNA and surface protein signals. The second dataset is obtained from an E18 mouse brain 9k cells generated on the Chromium scRNA-seq platform (10xGenomics) [11]. This dataset will cover some new cell types that are defined only in the upgraded model. We use this dataset to test the robustness of the expansion of cell-type catalogue. The third dataset was a 2.8k-tumor-tissue-associated-cells dataset obtained from a malignant pancreatic tumor tissue that was extracted from a genetically modified mouse (the KPC model) [12] generated on the Chromium platform. The study of the third dataset represents a typical scenario in which researchers seek to extensively interrogate a heterogeneous tissue sample using the scRNA-seq technique. The populations of the UC clusters and the SC predictions on these three datasets were compared. As it can be observed, some of the unsupervised clusters can be explained by SuperCT predictions, though some of the clusters might need further validation (**Supplementary Fig. 2**, **Fig. 3d, 3e, 3f**, **Fig. 4a and 4b**).

**Figure 2:**
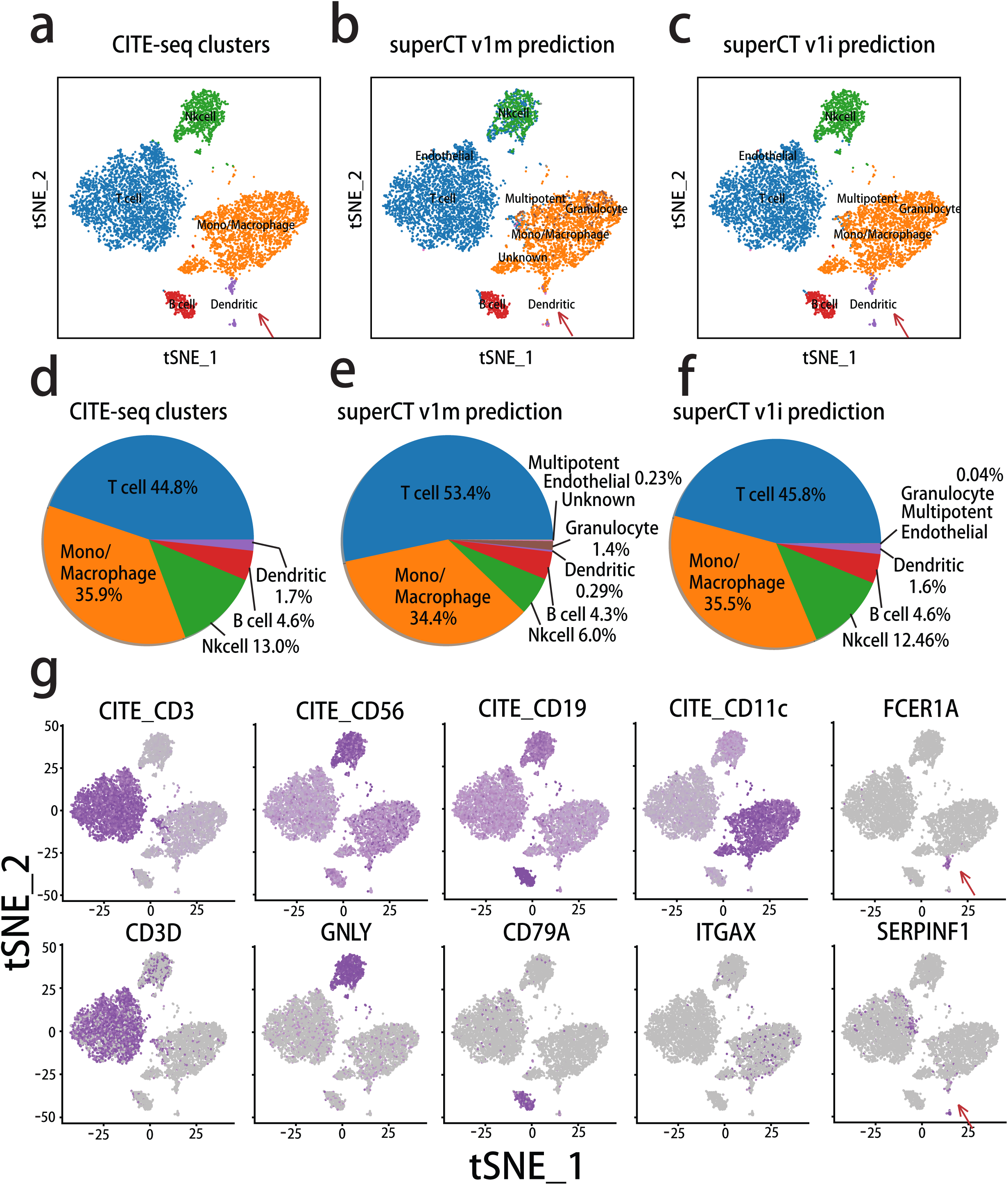
The upgraded predictions of CITE-seq human PBMC cells that are confirmed by the marker signals of both surface proteins and RNAs. a) The original cluster labels on the tSNE layout of CITE-seq PBMC cells. b) The prediction by SuperCT v1m. c) The prediction by SuperCT v1i. d) The Pie chart by the original CITE-seq PBMC cell types. e) The pie chart of the SuperCT v1m predictions. f) The pie chart of SuperCT v1i predictions. g) The signal distribution of the canonical marker genes for the PBMC subpopulations.

**Figure 3:**
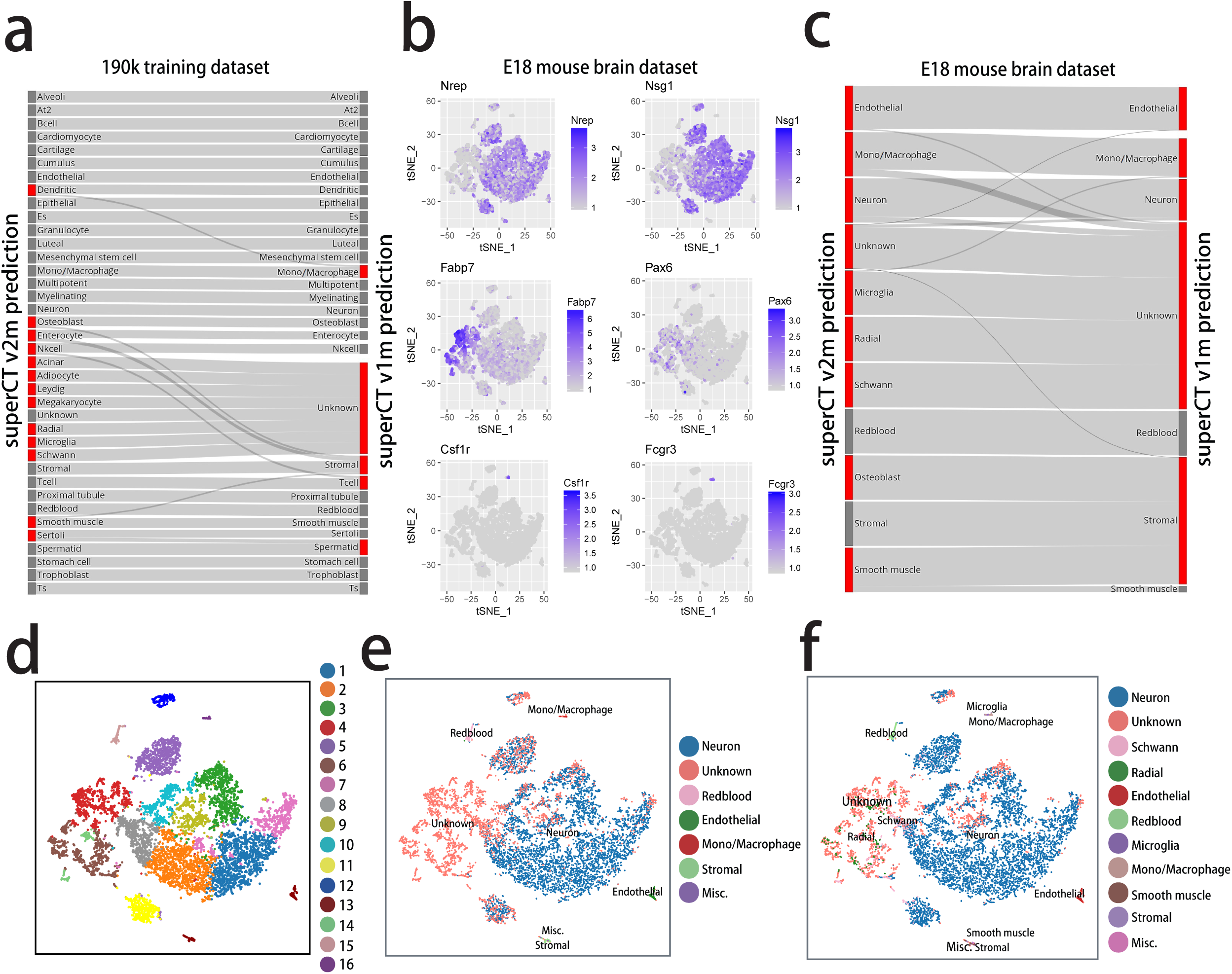
The assessment of the v1m-to-v2m upgrade by comparing the SuperCT predictions of different version in both 190k training dataset and E18 mouse brain dataset. a) The Sankey diagram indicating the upgrade of SuperCT v1i (right) labels from SuperCT v1m in 190k training dataset. b) The signal of the signature genes indicating the corresponding cell types of different clusters in E18 brain dataset. c) The Sankey diagram indicating the upgrade of SuperCT v1i (right) labels from SuperCT v1m (left) in the 9k E18 brain dataset. Most of the new cell types are from ‘unknown’ types. Some of the Microglia are from mono./macrophage type and some are from ‘unknown’ category. d) Unsupervised clusters of 9k E18 brain dataset e) SuperCT v1m predictions of 9k E18 brain dataset. f) SuperCT v2m predictions of 9k E18 brain dataset.

**Figure 4:**
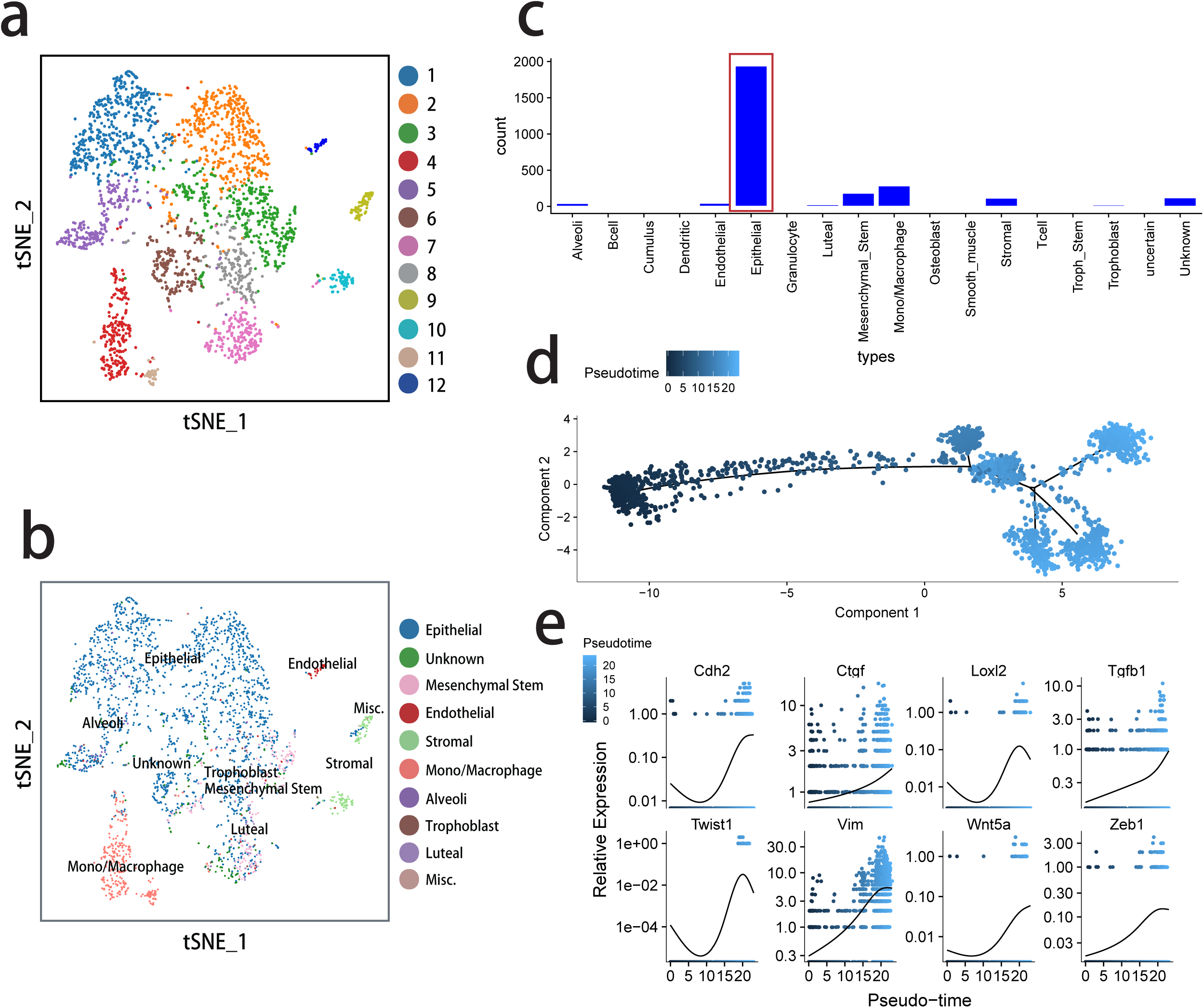
De-convolution of the signal of the constituent populations of mouse pancreatic tumor tissue. a) The unsupervised cell clusters of the mPDAC tissue defined by Seurat FindClusters function. b) The predicted cell types of the mPDAC tissue. The minor types with less than 5 cells are not shown in color legend. c) The bar-plot for the cells of the predicted cell types in mPDAC. The epithelial tumor cells in the red box are sorted out for the downstream analyses d) the pseudo-temporal trajectory of the tumor progression based on sorted tumor cells plotted by Monocle e) The EMT genes display up-regulations and correlate to the tumor progression.

There are three versions of classifier discussed in this paper, v1m, v2m and v1i. v1m and v1i both have 30 ‘known’ and one ‘unknown’ cell type in their catalogues. v2m has 37 ‘known’ and one ‘unknown’ type. ‘m’ stands for mouse and ‘i’ stands for immunity. Version number increase stands for the upgrade of the cell-type catalogue. **Fig. 1b** illustrates the two upgrade protocols that underlie typical evolution of SuperCT model. v1m is considered as the initial model trained from a large training dataset possibly with some error or insufficiency. In the v1m-to-v1i upgrade, we aim to test whether the online parameter optimization using additional training dataset from the similar background will make the classifier perform better in the target sample. In the v1m-to-v2m upgrade, we aim to demonstrate the robust expansion of the capability to predict more cell types with little sacrifice on the concordance over the predictions of other cell types.

We carefully examined the v1m predictions in reference to the original MCA cluster annotations using the confusion matrix across the defined cell types. The true predictions go to the diagonal of the matrix. Other than the undefined 7 cell types that go to ‘Unknown’ category, it is found that a few known cell types show lower concordance than the others (see confusion matrix in **Supplementary Fig. 1a**). Low concordance suggests the unsatisfactory performance of discerning these specific known types in the v1m training, most of which were difficult to annotate at the first place because of many shared markers. Even though the erroneous labels in MCA will compromise the training effect, it was still difficult to correct the original annotation without additional information to reach the ground truth in such large scale of data as MCA. Besides, in a testing dataset from human PBMC (CITE-seq), we found that v1m prediction result is at low concordance (88.3%), which is yet to improve (**Fig. 2a**). Other than the possible erroneous labels in the training dataset, the subtle discrepancy of the molecular signatures of immune cell types between mouse and human may also explains the low performance. Altogether, we believe the online optimization using higher confidence training data is necessary and can fix the problem. Given the availability of the datasets for a few of these types from another public resource [11], we were able to input more human immune cells from the tissue with higher confidence on the identity of human immune cells (peripheral blood) and thus to enhance the feature learning. This one is called v1m-to-v1i upgrade.

In the model update, instead of redoing the entire MCA 190k dataset in combination with the new PBMC 9k dataset, we implemented a more efficient ‘online learning’ strategy in the v1m-to-v1i upgrade. In online learning, ‘Catastrophic Forgetting’ is a tendency of ANN that the new training could make the model lose the ‘memory’ of the old training. This issue has been also taken care of using a ‘review training’ mechanism. The details for both strategies are described in the Online Methods. We used the CITE-seq PBMC dataset whose background is similar to the feature enhancement training dataset to assess the outperformance of v1i over v1m whose result is shown in **Fig. 2** and explained later.

In v1m-to-v2m upgrade, we specifically incorporated the ‘transfer learning’ strategy, which means some of the model parameters can be re-used in the new categorization tasks of the expanded reference catalogue. It is believed such strategy can improve the training efficiency by transfer knowledge from previous learning tasks [14]. More details are given in the Online Methods. The high concordance between v2m predictions to the original MCA annotation shown in following results suggests the success of transfer learning in cell-typing task.

In the quick assessment of the SuperCT v2m performance, **Fig. 1d** shows the predictions of SuperCT v2m colored by the cell types, which share the same color-code as original MCA cell-type annotation displayed in **Fig. 1c**. The overall similar color scheme suggests the high concordance. For the 190k MCA dataset, SuperCT v1m achieves 97.3% (171,677 cells) of concordance to the original labels in the 30 ‘known’ type category (176,675 cells). SuperCT v2m achieves 96.6% (174,739 cells) of concordance in the 37 ‘known’ type category (180,888 cells). Although the concordance rate seems a little bit lower in the v2m, total 3,062 more cells’ identities were accurately revealed (from unknown to specific known type) in the same dataset by the v2m. Both v1m and v2m give 100% concordant prediction to the MCA cells in the ‘Unknown’ category. In the mean time, Sankey diagram (**Fig. 3a and 3c**) shows this upgrade has limited influence on the predictions of other known types of v1m, which suggest the learning of the new cell types doesn’t make the model ‘confused’ over the other ones. In conclusion, the v2m does accurately characterize more cell types, thus outperforms v1m.

We were also able to rank the information gain of each gene’s presence/absence in discerning a cell type from the other types (The details are given in the Online Methods). The top 50 genes for each of 37 cell types are shown as examples in the **Supplementary Table 2.** These top-rank genes provide the guidance to discover the unique molecular features that underlie the v2m 37-cell-type categorization. Some of the top-rank genes are familiar to us, though many others have never been reported previously.

For the CITE-seq PBMC dataset, **Fig. 2a, 2b and 2c** show the layout of 4 predominant clusters, one minor population hidden at the lower corner (classical dendritic cells) and one minor segregated cluster (plasmacytoid dendritic cells). The original paper defined these 5 cell clusters based on their surface marker signals and the signature RNA signals. **Fig. 2b** shows the characterizations by SuperCT v1m. The overall concordance is 88.3%. **Fig. 2c** shows the characterizations by SuperCT v1i. The overall concordance is 98.5%. Fig 2d, 2e and 2f show the percentage of the annotated cell types. **Fig. 2g** shows more details of the marker signal of both surface proteins (top row) and RNA transcripts (bottom row) for the 5 cell types. The result suggests v1i gives the predictions with higher concordance than v1m in CITE-seq PBMC dataset.

In order to specifically validate the expandability of this SuperCT framework, we carefully examined in the change of the prediction result before and after the model upgrade in MCA training dataset and E18 mouse brain dataset. For the 190k training dataset, the Sankey diagram, **Fig. 3a** shows that the 7 new cell types defined by v2m were mainly derived from the ‘unknown’ types as defined by v1m. In E18 mouse brain dataset, 3 out of 7 additional cell types, (Radial glia, Microglia, and Schwann cells) in v2m which are nervous-system-related were anticipated in the mouse brain tissue. The Sankey diagram for this dataset, shown as **Fig. 3c**, provides the details on how the SuperCT v2m have learned the 3 new cell types from the ‘unknown’ classes or the similar cell types (microglia vs. mono./macrophage) from v1m. This result suggests, by increasing the output nodes of ANN model and including sufficient new training cells, SuperCT is able to predict more specific cell types. Interestingly, v1m made an ‘excusable’ mistake by taking microglia cells as monocyte/macrophages in v1m (**Fig. 3a, 3c and 3e**). This ‘mistake’ can be explained by the similar roles of microglia to the nervous system as the macrophage to the part of a mammalian body. **Fig. 3b** and **Supplementary Table 2** also shows multiple genes, such as colony-stimulating factor receptor (CSF1R) and Fc fragment of IgG receptor (FCGR3A), are shared by mono./macrophage and microglia cells. **Fig. 3e and 3f** show the tSNE layout of the predicted cell types by v1m to v2m, suggesting the successful evolution of the model predictions from unknown to specific cell types that gradually uncover the role of the unsupervised cell clusters in the **Fig. 3d**.

At last, we summarize the SuperCT result of a pancreatic ductal adenocarcinoma dataset derived from KPC mouse (mPDAC, dataset #3). **Fig. 4a** shows Seurat FindClusters function yielded 12 putative cell clusters while SuperCT characterized total 17 cell types (**Fig. 4c**). We were particularly interested in the primary epithelial tumor cells, which had been assigned to ‘Epithelial’ type by SuperCT v2m (**Fig. 4b** and **4c**). We then perform the *in silico* ‘sorting’ of the predicted 1940 epithelial tumor cells. The typical mPDAC markers, e.g. EPCAM, cytokeratin 19, CD133 etc., were indeed enriched in these cells, validating the tumor cell identity (**Supplementary Fig. 3**). With the cell identity information brought by SuperCT v2m, we were able to further explore whether any dynamic change of the molecular signals could be observed in the predicted epithelial tumor lineage. The pseudo-temporal analysis was performed on these ‘sorted’ tumor cells using Monocle 2 method [13]. A tumor progression trajectory was thus derived. The order information of the tumor cells in the inferred trajectory was utilized to perform the regression analysis in order to reveal the relationships between the tumor progression and the gene expression change. **Supplementary Table 3** shows a list of genes that give significant p-values for the regression coefficient. (See details in the Online Methods). A number of the correlated genes, such as N-cadherin, Twist1, Loxl2 etc., that are involved to the epithelial-mesenchymal-transition (EMT) were identified. (shown in **Fig. 4e**) Other than the epithelial tumor population we have extensively discussed as above, the clusters C4+C11, C9+C10 and C12 defined by Seurat FinderClusters function are corresponding to the SuperCT predictions of monocyte/macrophage, stromal and endothelial cells (**Fig. 4a** and **4b**). In here, SuperCT gives explicit interpretation on these clusters.

It is notable, the MCA dataset was generated on the Microwell-seq platform and the 4.2k PBMC dataset was generated on the Chromium scRNA-seq platform (10xGenomics). Three testing datasets are generated on the platforms of CITE-seq, and Chromium scRNA-seq platforms, respectively. All these platforms produce the typical digital expression matrices based on Unique-Molecular-Index (UMI) counts. Moreover, CITE-seq PBMC dataset was generated from human tissue whereas the MCA, E18 brain and mPDAC datasets were generated from mouse tissues. The reliable results suggest the compatibility of SuperCT framework is not only across different scRNA-seq platform but also across different mammalian species, though the online optimization like v1m-to-v1i upgrade would benefit the application of the specific scenario.

## Discussion

In this study, using training and independent testing datasets, we have demonstrated the reliable performance of the supervised-learning-based classifiers in the characterization of the previously defined cell types in the heterogeneous tissue samples. This is the first scRNA-seq analytical framework that is independent of unsupervised clustering algorithm. With least requirement of expert-knowledge or bioinformatics skillset from the individual user, the SuperCT classifier delivers accurate cell types information for thousands of single-cells in just a few minutes.

A classifier with good performance is not built overnight. Neither is it a one-size-fit-all model. It is actually the growing human learning that drives the machine learning, which means a continual learning mechanism in this framework should be designed at the very beginning. The two successful upgrades described in this paper have proved that the online-learning and the transfer-learning algorithm incorporated in the SuperCT framework make the model evolution not only possible but efficient.

In general, supervised and unsupervised learning methods play the complementary roles in the growing findings facilitated by machine. The classification results from both methods in the analyses of scRNA-seq data can be the cross-reference to each other. As it has been seen, many of the cell grouping defined by UC can be validated by SC in many cases. The cell grouping by SC takes the full advantage of the *a priori* learning that makes the cell type prediction less subjective but more efficient. The UC cell grouping is driven by the current data, whose new information could lead to the new findings. Three possible scenarios in UC and SC comparison should be further explored in order to achieve new findings. 1) The cells form segregated clusters by a UC method with cluster-specific molecular features but SC characterization may give only one cell-type. 2) There is an obvious cluster, which may be categorized as ‘unknown’ by SC. 3) There is a cell type that is characterized in the tissue that may ‘not make any sense’, such as the ‘monocytes’ were found in mouse brain by SuperCT v1m.

## Availability

The webapp for different versions of SuperCT can be accessible from the following URL. https://sct.lifegen.com/.

### Contribution

PX, MZ and WL conceived the idea. PX and MG implemented the algorithm and performed analyses. MG prepared most of the illustrations of the manuscript. CW and PN provided the bench-side support of scRNA-seq data generation. CY, HH and DVH provided valuable suggestion on the application and editing on the manuscript. MZ and WL prepared the manuscript.

## Online Methods

### Implementation of the SuperCT Artificial Neural Network

The neural-network structures, the learning algorithms are implemented using Keras API. We design a fully connected artificial neural network (ANN) model for the cell type classification. The inputs are the binary signals of 16,013 genes that are homologous between human and mouse. We include these homologous genes to adapt the model to the application of both human and mouse study in this paper. To enhance the compatibility across different scRNA-seq platforms, we convert the digital expression values to the binary values, which means the genes are either present or absent in the cells. The input layer is connected with a hidden dense layer with 200 neurons and the first layer is fully connected to the next 100 neurons, respectively, using ReLU (Rectified Linear Unit) activation functions. Two random neuron dropouts (dropout rate at 0.4) occur after each layer in order to control over-fitting. The number of the output nodes corresponds to the number of the cell types in the catalog, which is 30+1 for v1m/v1i and 37+1 for v2m respectively. As the sample sizes of the different cell types vary from hundreds to tens of thousands in MCA dataset, to avoid the under-representation of the small-sample-size cell types in the calculation of the accuracy function, we include the class-weight based on the sample size of each type in the model training. The loss function is defined as categorical cross-entropy.

### Organization and preprocessing of the training dataset

There are three versions of SuperCT, v1m (‘m’ stands for mouse), v1i (‘i’ stands for immune cells) and v2m. We firstly select a total of 176,675 cells in the 30 categories of known cell types defined in mouse cell atlas project (MCA). These 30 cell types have more than 1000 counts in MCA. We then selected 4,227 cells (4.2k-new-type-cells) in the 7 categories of known cell types other than the 30 types in MCA. These 7 cell types have more than 500 counts in MCA. We also synthesized 8,923 scRNA-seq profiles (9k-synthetic-cells) by shuffling the randomly selected MCA scRNA-seq profiles. Total 189,825 labeled single-cell expression profiles (refer to as the 190k training dataset) were used for the training of SuperCT v1m and v2m. To test the robustness of the model expansion from v1m to v2m, the variable training labels of the two versions are described as follow. Other than the 9k-synthetic-cells that are always assigned to ‘unknown’ category, the 4.2k-new-type-cells were assigned to ‘unknown’ category in v1m but assigned to 7 specific types in v2m. We are wondering whether the extra 7 types could be learned from the ‘unknown’ category in the new training. Other than the abovementioned MCA and synthesized cells, 8,976 additional PBMC cells (9k-immune-cells) from 10xGenomics public database are included for SuperCT v1i training. In order to make fair comparison, the 4.2k-new-type-cells were still assigned to the unknown category in v1i because v1i is considered as the optimization of v1m.

### Initial training and continual learning of the models

For the SuperCT v1m, we use the batch learning of the entire 190k MCA data. We applied SGD (Stochastic Gradient Descent) by dividing training data into mini-batches of size 1,024. Numbers of training iterations were manually determined based on the gap between the training and validation error to control over-fitting (early stopping). In the two upgrades from v1m to v1i and v1m to v2m, we took the v1m parameters to initiate the online learning. We also took the mini-batches of 1,024 from the new training data and update the model parameters. In order to alleviate the ‘Catastrophic Forgetting’, we designed a ‘review training’ mechanism to ‘refresh’ the memory of the previous learning in the v1m. Randomly selected 4000 cells from the previous training dataset are fed again for the model training after each epoch of learning by the new training dataset.

### Transfer learning in the model expansion

In the training of the new cell types using MCA dataset, we freeze the first hidden layer (200 nodes) and only update the weights of the second layers and the output layer. This learning strategy will retain the transformed features residing in the hidden layer in previous training and use them for the characterization of more cell types.

### scRNA-seq dataset generated from mouse pancreatic tumor tissue

Freshly harvested tumors from KPC (LSL-KrasG12D/+; LSL-Trp53R172H/+; Pdx-1-Cre) mice were subjected to mechanical and enzymatic dissociation using a Miltenyi gentleMACS Tissue Dissociator to obtain single cells. The 10X Genomics Chromium Single Cell 3′ Solution was employed for capture, amplification and labeling of mRNA from single cells and for scRNA-Seq library preparation. Sequencing of libraries was performed on an Illumina HiSeq 2500 system. Sequencing data (fastq files) was input into the CellRanger pipeline to align reads and generate gene-cell digital expression matrices.

### Unsupervised clustering, dimensional reduction and data visualization

Most of the unsupervised clustering, dimensional reduction and data visualization in this paper are accomplished by a commonly used scRNA-seq analytical suite, Seurat. The Seurat objects are generated for each dataset with their digital expression matrices as input. The PCA is performed by Seurat RunPCA function. The tSNE coordinates are calculated using Seurat RunTSNE function. The putative clusters are defined by Seurat FindClusters function using the top 10 principle components and other default parameters. The prediction result are loaded to the Seurat metadata in order to be shown in reference to the tSNE layout.

### Top signature genes for each cell type ranked by information gain

In order to calculate and rank the information entropy gain, the following equations are used.

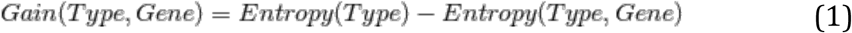

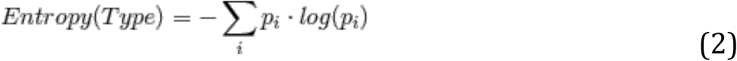

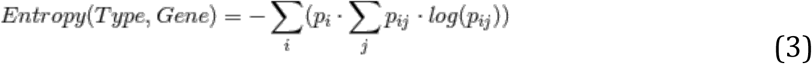

The binary values of each gene over the binary values as a certain type for the total cells are utilized to make a contingency table in order to calculate the p in Eq. 2 and Eq. 3. Whereas pi denotes the priori probability of cell belonging to the cell type i; p_ij_ denotes the probability of cell belonging to the cell type i (yes or no) when gene signal is in j status (present or obsent). The top 50 genes are ranked based on the value of information gain (Eq. 1) in **Supplementary Table 2** for each type.

### Search of genes that correlate to the tumor progression

The epithelial tumor cells are characterized from the scRNA-seq data of the mouse PDAC tumor tissues using SuperCT v2m. These cells’ digital expression profiles were pulled out as the input to calculate the pseudo-time ordering using Monocle2. The pseudo-time values of each cell and the gene expression value for that cell were used to fit a linear model using an R function ‘lm’. The p value of the coefficient of each linear model is used to determine whether the gene expression correlates to the tumor progression by time.

## References

1. Shekhar, K. et al. Cell 166, 1308-1323.e1330 (2016).

2. Tasic, B. et al. Nat. Neurosci. 19, 335–346 (2016).

3. Zeisel, A. et al. Science 347, 1138–1142 (2015).

4. Yao, Z. et al. Cell Stem Cell 20, 120–134 (2017).

5. Soares, J.V.B., Leandro, J.J.G., Cesar, R.M., Jelinek, H.F. & Cree, M.J. IEEE Transactions on Medical Imaging 25, 1214–1222 (2006).

6. Geng, X., Zhan, D.C. & Zhou, Z.H. IEEE Transactions on Systems Man and Cybernetics Part B-Cybernetics 35, 1098–1107 (2005).

7. Camps-Valls, G., Bandos, T.V. & Zhou, D.Y. IEEE Transactions on Geoscience And Remote Sensing 45, 3044–3054 (2007).

8. Rämö, P et. al. Bioinformatics 25(22), 3028–3030 (2009)

9. Han, X. et al. Cell 172, 1091-1107.e1017 (2018).

10. Stoeckius, M. Nature Methods 14, 865–868 (2017)

11. https://www.10xgenomics.com/solutions/single-cell/

12. Olive, K.P. et al. Science 324, 1457–1461 (2009).

13. Qiu, X. et al. Nature Methods 14, 979–982 (2017)

14. Thrun, S., Advances in Neural Information Processing Systems 8, 640–646 (1996)

